# Specific Peptides Predict Protein Classification

**DOI:** 10.1101/2022.02.04.479085

**Authors:** David Horn, Uri Weingart

## Abstract

The methodology of Specific Peptides (SP) has been introduced within the context of enzymes. It is based on an unsupervised machine leaning (ML) tool for motif extraction, followed by supervised annotation of the motifs. In the case of enzymes, the classifier is the Enzyme Classification (EC) number. Here we demonstrate that this method reaches precision of 96.5% and recall of 89.1% on presently available protein sequences. We also apply this method to two other protein families, GPCR and ZF, find their corresponding SPs, and provide the code for searching any protein sequence for its classification under any such family.

## 1. Introduction

Genes were perceived well before they have been determined to exist on chromosomes. In hindsight, it seems quite a surprise to find that they are just stretches of nucleotides within much larger sequences of DNA, often also interspersed by non-coding sections (introns). The identity of genes comes to life after being transcribed into RNA molecules, and translated into proteins, the important components of the machinery of living cells.

Proteins are molecular chains of amino acids. They are being studied by investigating the linear composition of amino-acid sequences, or their folding structures, or their functional properties, as revealed by their interactions with other molecules. In this paper, we discuss a different perspective of their structures, resulting from amino acid motifs, which are observed to be common to many proteins having the same function, or belonging to homolog genes of different species.

We follow the methodology developed and tested in [1-3], pointing out the existence of **Specific Peptides (SP**s) which are motifs of length ≥ 7 amino acids, occurring on enzymes only. We reanalyze all enzymes using the updated Enzyme Classification (EC) labelling, employed by SwissProt [4]. This analysis demonstrates the high predictive power of enzymatic SPs which will be labelled ESPs. This is followed by analyzing GPCR proteins, including the large group of non-OR proteins, expanding the results of a previous study of OR proteins [5]. We then continue to analyze Zinc Finger proteins and find their relevant SPs. Thus we end up with SPs defined for proteins belonging to all these families.

The analysis starts with the motif extraction method MEX [6], which is an unsupervised algorithm finding motifs with high occurrence in a given text. In enzyme classification this text contains 90% of all enzymes in the data set [4], to be labelled as the training set Ptrain. Next we use a supervised methodology: labelling all EC motifs according to the EC assignments of proteins in Ptrain, and discarding motifs which appear in Ntrain, which contains 90% of the non-enzymatic proteins in the data. The prediction accuracy is finally tested on the remaining 10% of the data, Ptest and Ntest.

Clearly assignments of SPs should be considered within the context in which they were derived. They are not supposed to annotate a free peptide, but only the motif appearing within a protein sequence. Still, as such, they can help identifying and annotating novel proteins, and may turn out to be very useful for artificial protein engineering [7] and for medical research and development [8].

## 2. Results

### 2.1 Enzyme Specific Peptides

The SwissProt entry [4] (version 2021_01) contains 564,227 proteins of many species. In order to enable a training and testing procedure we divided randomly the enzymes which had a single EC annotation into two sets: 227,488 were designated to a training set (Ptrain) and 25,309 enzymes were designated to a test set (Ptest). In parallel we also constructed non-enzymatic training and test sets, Ntrain and Ntest, containing 264,739 and 29,416 proteins correspondingly. This validation set serves to discard motifs which are not specific to enzymes.

Using the Enzyme Classification (EC) nomenclature, enzymes are classified into seven classes, EC1 to EC7, and within each EC class they are grouped into a hierarchy of four levels.

Some are classified just into the first level, numbered by the class, some at levels 2 or 3, but most at level 4, which is often associated with homologs of the same gene in different species. Proteins which have enzymatic regions belonging to two different EC classes were discarded from the training set.

Following [4] we restricted our MEX search to motifs of length ≥7 amino acids. Details of our procedure of analysis are explained in the Methods section. Our procedure leads to a set of 286,755 specific peptides which we label as ESPs. They are provided as a Json list in our github entry [9] which also includes the code for searching a protein for the occurrence of such ESPs.

In order to test the usefulness of ESPs in predicting the EC labelling of a protein, we ran it on the test sets Ptest and Ntest. An SP hit on Ptest is regarded as true positive (TP) if the Swissprot EC assignment of the enzyme appears on the EC tree of the SP. If no SP hits an enzyme, it is labelled as false negative (FN). If an SP hits a protein in Ntest, the latter is declared as false positive (FP). If no SP hits a protein in Ntest, it is regarded as true negative (TN).

The results are presented in Table 1:

**Table 1.** Classification of Enzymes according to ESPs.

| TP | FP | FN | TN | Precision | Recall |
| --- | --- | --- | --- | --- | --- |
| 22,283 | 806 | 2,726 | 28,910 | 96.5% | 89.1% |

We use conventional definitions of Precision=TP/(TP+FP) and Recall=TP/(TP+FN). The sizes of Ptest and Ntest are 25,309 and 29,416 correspondingly. The 806 FP events include 300 from Ptest (with mismatched EC assignments) and 506 from Ntest.

### 2.2 GPCR

G protein coupling receptors (GPCR) play dominant roles in olfaction, vision and many other cellular functions.

Olfactory Receptors (OR) were studied in [5] using motifs of length ≥5 derived by the MEX methodology. They [5] have demonstrated how the resulting motifs can be employed in providing the sketch of an evolutionary tree of species, and have provided a web-service for OR protein assignment on the basis of these motifs.

We extend our analysis to all Swissprot GPCRs. After motif extraction we start with human GPCRs, and exhibit the specificity of all motifs of length ≥ 7 to either OR proteins, or to non-OR (NOR) proteins within all GPCRs. There exist 156 OR motifs of length ≥ 7 with hits on the 469 human ORs, and 2896 NOR motifs hitting the 148 human NOR proteins. There is no overlap between these lists, i.e. they are specific to either OR or NOR proteins.

While the number of human OR proteins (469) is larger than the NOR proteins (148), the number of the NOR motifs is much larger (2896 vs 156 of length ≥ 7) because of the large variety of modalities which are served by NOR proteins. Turning to all Swissprot GPCRs, for all organisms, we expect to find a clear separation between OR and NOR, as well as discover a very large number of protein-SP biclusters in the NOR GPCRs. We find 562 OR proteins, leading to 367 OR motifs with length ≥ 7, and 2481 NOR proteins with 3710 corresponding motifs. Once again, the two sets of motifs are specific to the two sets of proteins. We then proceed to search for protein-SP biclustering of the NOR data. The large number of NOR proteins and motifs allows for their sorting into clusters, as listed in Table 2. Both OR and NOR SPs will be referred to as GSPs. Their lists are provided as Json files in our github entry [9].

**Table 2.**
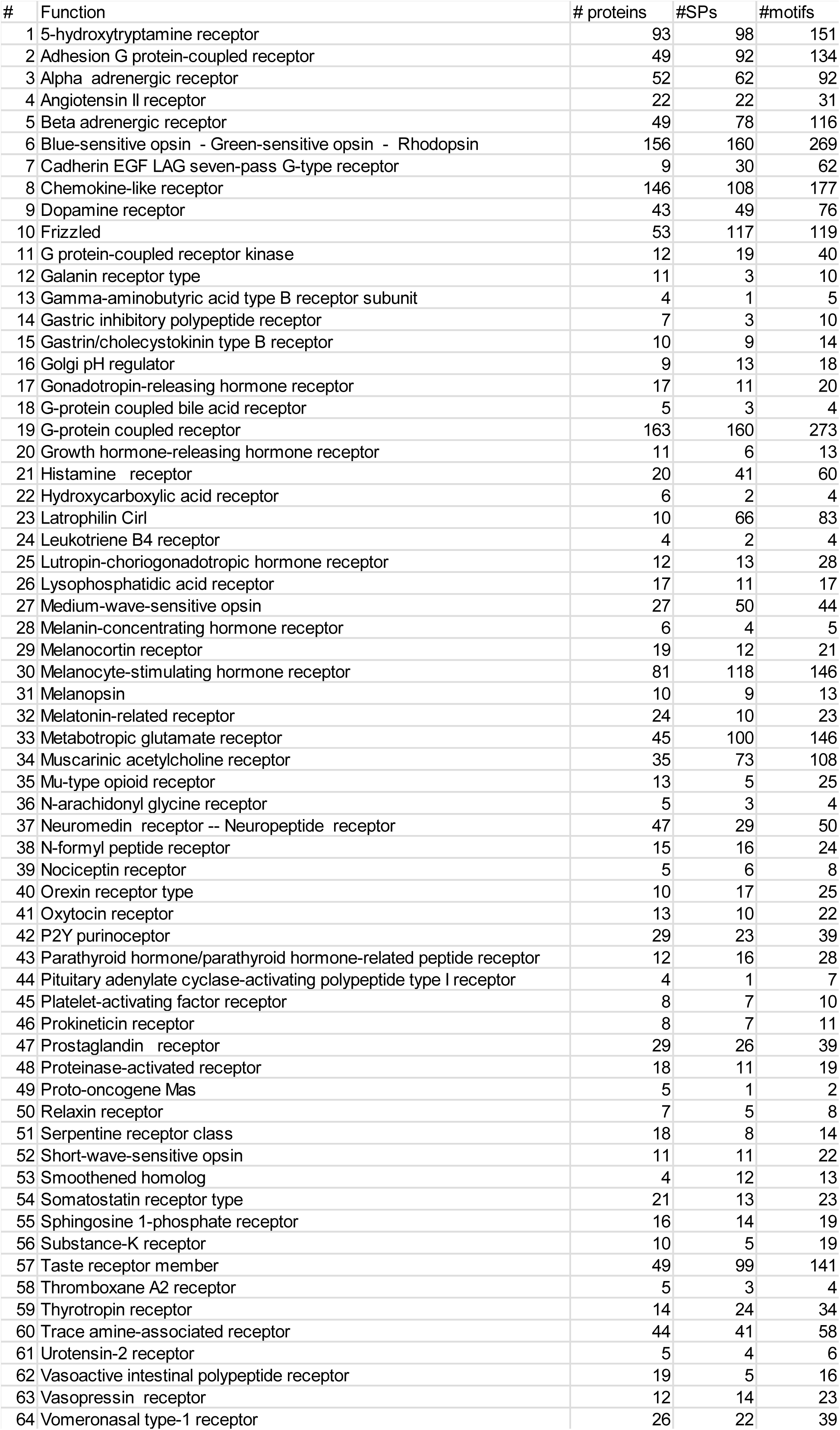
64 Protein clusters among NOR GPCR. #SPs refers to those which are specific to the cluster, while #motifs refers to other GSPs occurring in other NOR clusters as well.

Next we run all the GSPs against the Enzymes of our EC study. We find 63 hits of these motifs on 3 EC classes, thus providing EC identifications of 3 NOR classes. They are listed in Table 3:

**Table 3.** Three NOR classes which belong to three EC classes.

| EC classification | NOR classification |
| --- | --- |
| 2.7.11.14 Rhodopsin kinase | Blue-sensitive opsin, Green-sensitive opsin, Rhodopsin |
| 2.7.11.15 [Beta-adrenergic-receptor] kinase | Beta adrenergic receptor |
| 2.7.11.16 [G-protein-coupled receptor] kinase | G protein-coupled receptor kinase |

There exist 20 other sporadic hits of NOR GSPs on EC proteins, which are consistent with expected noise.

### 2.3 Zinc Finger proteins

We have analyzed 2582 Swissprot ZF proteins and extracted 1487 motifs of length ≥7 which are declared to be ZSPs. 786 of all the proteins are human ZF proteins, and they display hits by 1412 of the SPs.

Since ZF proteins may contain several ZF domains, we encounter reappearance of motifs on different locations within the same protein. This is different from our previous studies of EC and GPCR proteins, where inter-protein multiple appearances were responsible for the generation of MEX motifs. Here we find that intra-protein recurrences play an important role.

To illustrate this fact, we display in Table 4 some of the “popular” ZSPs, which have 100 or more hits on all human ZF proteins. On the right of the table we provide the sum of all shown hits for each protein as displayed here, and compare it to the total number of ZSPs hitting each protein. On the bottom we provide analogous counts for each ZSP.

**Table 4.**
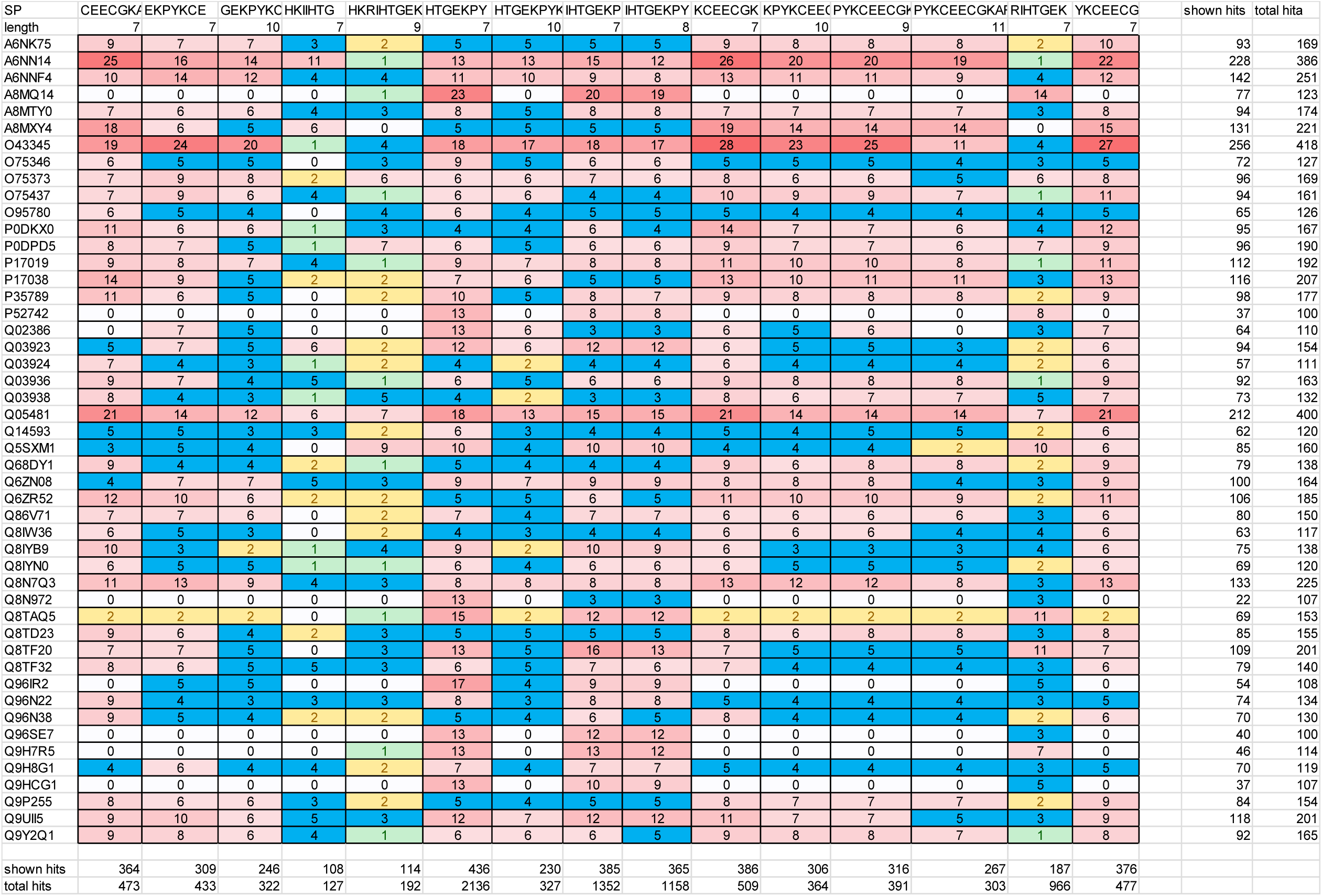
Number of hits by different SPs, displayed on different human ZF proteins. Large numbers correlate with the fact that many ZF regions can be found on the same protein.

It should be realized that SPs of length n can be contained within SPs of length >n, as can be seen in this table, which serves as an examples rather than a summary. Summary of all ZSPs and their hits on ZF proteins is provided in our github entry [9].

Clearly the repetitive appearances of SPs on a given protein reflect the existence of many ZF regions on the same protein. The latter is usually larger than the number of repeats of a single SP, since different SPs may belong to different ZF regions.

There exist some proteins which act as enzymes and possess zing fingers. One outstanding example is PRDM9. This protein serves recombination hotspots during meiosis by binding nucleotides with its zinc fingers. The annotations of the human version of this protein are provided by https://www.uniprot.org/uniprot/Q9NQV7. They contains 14 ZF regions. The first starts at location 388 and has length of 24 amino acids. The other 13 start at 524 and are of length 23 each. In Fig. 1 we display, in color code, the loci of hits by all ZSP and ESP motifs of length ≥ 7.

**Fig. 1.**
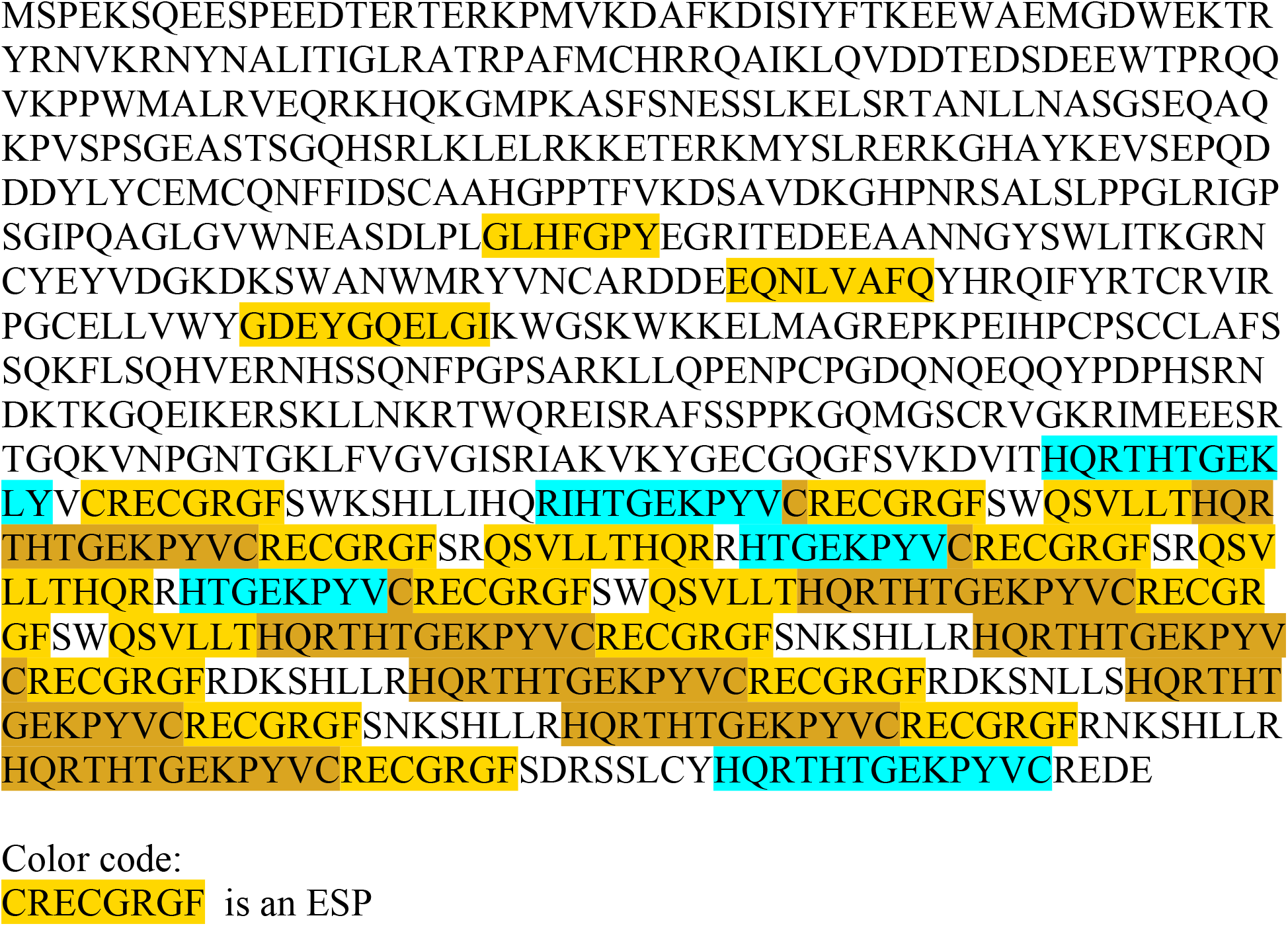
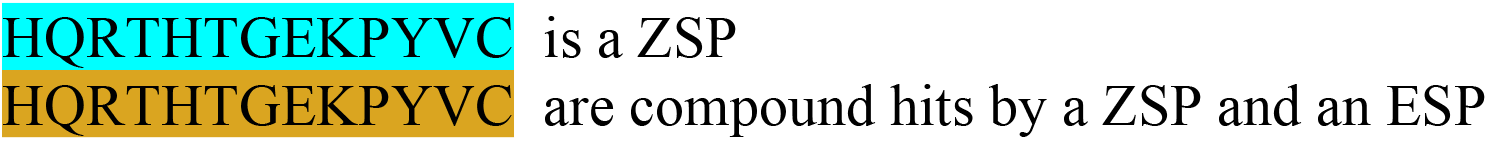
The sequence of PRDM9_HUMAN Histone-lysine N-methyltransferase (Q9NQV7) and color coded display of hits by ESPs of EC 2.1.1.43 and ZSPs, which may partially overlap each other.

Fig. 1 shows that the last 12 ZF domains display very high similarity and an exact periodicity. Comparing with Uniprot annotations we find that all ZF domains have the structure YVCRECxxxxxxxxHQRTHT, where the additional 8 amino acids, replaced by x, vary according to the nucleotide targeted by the ZF domain. In Fig. 1 we note prevalent occurrence of the structure HQRTHTGEKPYVCRECGRGF which includes the suffix of a previous ZF domain and the prefix of the next ZF domain. The colors reflect occurrences of hits by ZSPs and ESPs. Between the suffix of one structure and the prefix of the next we find the quintet GEKPY which fills the gap between 23, the length of the ZF motif, and 28, the regular periodicity observed in this protein over the range of its last 12 ZF regions.

## 3. Summary and Discussion

Our methodology is based on machine learning (ML) practices: MEX is an unsupervised tool for motif extraction; these motifs are then searched on protein sequences using supervised annotation to classify the results. In the case of enzymes, the classifier is the Enzyme Classification which is defined in terms of seven classes and four levels in each class.

ESPs are specific peptides whose presence on the amino acid sequence of the protein indicates its EC number, as well as the tree associated with it. This methodology was introduced in 2007 [1]. Other ML studies appeared in the meantime, trying to solve the same (or related) problems using various ML tools. Many neglected to notice that SPs can do the required EC prediction quite well, often even better than the new tools.

Some examples of recent ML methodologies are DeepEC [10] and MAHOMES [11]. DeepEC employs 3 deep convolutional neural networks and a homology analysis tool to the study of enzyme sequences. When applying it to a test set which uses 201 enzymes they obtained precision = 0.92 and recall = 0.455 (quoted from Table 2 in [10]). This is considerably worse than our results in Table 1, which were based on a much larger (25K) test set. Other five ML methods which they [9] compared themselves to, were even worse.

MAHOMES[11] uses a decision-tree ML model, which is structure-based, employing physicochemical features specific to catalytic activity. Their main aim is to classify metals bound to proteins as enzymatic or non-enzymatic, and they succeed doing it with precision=0.922 and recall=0.901. Comparing to sequence-based technologies, they find that DeepEC scores on their tasks are precision = 0.905 and recall = 0.596. They find that another homology method, EFICAz2.5 [12] (which lost to DeepEC according to [10]), had better statistics (precision=0.922 and recall=0.901) but still falls short of their own [11]. For an older review of ML studies of enzymes see [13].

Our precision/recall results attest to the usefulness of the MEX unsupervised methodology in discovering relevant and unique motifs, the specific peptides (SPs). Our approach is not limited to enzyme studies. We have demonstrated this flexibility by investigating GPCR and Zinc-finger proteins, leading to a wealth of novel SPs. We provide in [9] a documented python code which allows for SP searches of all the functionalities which we have studied. It contains the lists of 2,002 NOR GSPs, 351 OR GSPs and 1,482 ZSPs in addition to the 286,755 ESPs.

## Methods

### Building the list of ESPs

In order to run the Motif Extraction program (MEX) [6], we divided the enzymes training set into batches grouped by joint level 2 assignments, and batches of enzymes with single level 1 assignments. Following [4] we restricted our MEX search to motifs of length ≥7 amino acids. The analysis led to 307,989 motifs. All motifs were then annotated after collecting the information of the IDs of enzymes hit by a particular motif (i.e. occurring in full on the amino acid chain of the enzyme) and how many times was a particular enzyme hit by a particular motif.

The EC number description, indicating both class and level, can be viewed as an inverted tree with a maximum depth of 4. For every motif, we map the EC numbers of the enzymes it hits on the training set onto a single EC tree. Starting from level 4 and moving upwards, we search for the first level which is a unique descendent of a higher level. The EC number of this unique descendant is assigned to the motif.

In order to remove motifs which may occur also on non-enzymatic proteins, we search for hits of all motifs on the non-enzymatic Ntrain set. Such motifs are removed from the list of specific peptides. Thus, to summarize, a motif of length ≥ 7 amino acids is labeled as an Enzyme Specific Peptide (ESP) if:

-it hits (i.e. appears in full on the amino acid chain of) enzymes belonging to only a single EC class of Ptrain

-and it does not hit any protein in Ntrain

This procedure leads to the reduction of the set of motifs to 286,755 specific peptides which we label as ESPs. They are provided as a Json list in our github entry [9] which also includes the code for searching the sequence of a protein for the occurrence of such ESPs on its amino acid chain..

